# A single shot of a hybrid hAdV5-based anti-COVID-19 vaccine induces a long-lasting immune response and broad coverage against VOC

**DOI:** 10.1101/2021.08.11.455942

**Authors:** M. Verónica López, Sabrina E. Vinzón, Eduardo G. A. Cafferata, Felipe J. Nuñez, Ariadna Soto, Maximiliano Sanchez-Lamas, Jimena Afonso, Diana Aguilar-Cortes, Gregorio D. Ríos, Juliana T. Maricato, Carla Torres-Braconi, Vanessa Barbosa-da Silveira, Tatiane Montes-de Andrade, Tatiana Carvalho-de Souza Bonetti, Luiz M. Ramos Janini, Manoel J. B. Castello Girão, Andrea S. Llera, Karina Gomez, Hugo H. Ortega, Paula M. Berguer, Osvaldo L. Podhajcer

**Affiliations:** Laboratory of Molecular and Cellular Therapy, Fundación Instituto Leloir – CONICET (IIBBA), Buenos Aires, Argentina; Laboratory of Immunology and Molecular Microbiology, Fundación Instituto Leloir – CONICET (IIBBA), Buenos Aires, Argentina; Securitas Biosciences, Montevideo, Uruguay; Fundación Instituto Leloir, Buenos Aires, Argentina; Discipline of Microbiology, Department of Microbiology, Immunology and Parasitology – DMIP, Paulista School of Medicine, Federal University of São Paulo – UNIFESP, Sao Pablo, Brazil; Discipline of Gynecology, Department of Medicine, Paulista School of Medicine, Federal University of São Paulo – UNIFESP, Sao Pablo, Brazil; Instituto de Investigaciones en Ingeniería Genética y Biología Molecular Dr. Héctor N. Torres (CONICET). Buenos Aires, Argentina; Centro de Medicina Comparada (ICiVet-Litoral), Universidad Nacional del Litoral – CONICET, Santa Fe, Argentina

## Abstract

Most approved vaccines against COVID-19 have to be administered in a prime/boost regimen. We engineered a novel vaccine based on a chimeric hAdV5 vector. The vaccine (named CoroVaxG.3) is based on three pillars: i) high expression of Spike to enhance its immunodominance by using a potent promoter and a mRNA stabilizer; ii) enhanced infection of muscle and dendritic cells by replacing the fiber knob domain of hAdV5 by hAdV3; iii) use of Spike stabilized in a prefusion conformation. Transduction with CoroVaxG.3 expressing Spike (D614G) dramatically enhanced Spike expression in human muscle cells, monocytes and dendritic cells compared to CoroVaxG.5 that expressed the native fiber knob domain. A single dose of CoroVaxG.3 induced potent humoral immunity with a balanced Th1/Th2 ratio and potent T-cell immunity, both lasting for at least 5 months. Sera from CoroVaxG.3 vaccinated mice was able to neutralize pseudoviruses expressing B.1 (wild type D614G), B.1.117 (alpha) and P.1 (gamma) Spikes, as well as an authentic WT and P.1 SARS-CoV-2 isolates. Neutralizing antibodies did not wane even after 5 months making this kind of vaccine a likely candidate to enter clinical trials

## INTRODUCTION

The disease caused by the novel coronavirus severe acute respiratory syndrome coronavirus 2 (SARS-CoV-2) has had an effect of enormous proportions, globally leading to more than 200 million confirmed cases and causing more than 4 million deaths (https://covid19.who.int/). During the last months different regulatory bodies issued emergency use authorization for different vaccines based mainly on mRNA and adenoviral-based platforms^1^. With the sole exception of the hAdV-26 based vaccine that was approved for one dose administration after showing an efficacy in clinical trials slightly over 65 %^2^, all the vaccines are given as a prime-boost approach to achieve maximal immune response and protection^3^. Companies are facing challenges in manufacturing vaccines and building the supply chains to meet the demand for COVID-19 vaccines. Different countries prioritized distribution of a first vaccine dose to as many people as possible^4^. Thus, the need for two doses and the fact that mRNA vaccines require logistically difficult cold-chains ^5^ make these type of vaccines more challenging to deploy in developing countries where ultra-low freezers may not be widely available. Therefore, getting a single dose vaccine with suitable stability and storage properties that can reach rapidly the local population is a major challenge for the scientific community especially in low and middle income countries.

Despite the unprecedented achievement of having approved vaccines in one year it is still early to establish the durability and extent of the protection, and recent data on the vaccines which first received approval, like BNT162b2 from Pfizer-BioNTech, point to a diminished efficacy already six months after vaccination^6,7^. Most importantly, it is still unclear the way to optimize the existing vaccines to protect against the prevalent SARS-CoV-2 variants of concern (VOC) that are spreading globally^8^. These VOC have multiple mutations in Spike, mainly in the RBM region and N-terminal domain, that differ substantially from the Spike variants encoded by the already approved vaccines. In fact, all the evidence indicates that serum samples obtained from convalescent people or from vaccines offer diminished protection against the β, γ and δ variants^9^. Replication incompetent adenoviral-based vaccines can induce a long term adaptive neutralizing humoral and cellular immunity^10^. The prevalence of anti-hAdV5 immunity in the developing world in the past^11^ led different academic groups and companies to identify less prevalent human adenovirus serotypes^2,12^ or non-human primates adenoviral vectors^13^ for anti-COVID-19 vaccine development. However, previous studies that compared the immunogenicity induced by hAdV5-based vaccines with the less prevalent human adenovirus hAdV26 and the chimpanzee-derived adenovirus ChAdOx1 among others, demonstrated that hAdV5 induced the most potent immune responses^10,14,15^. Moreover, hAdV5 is no longer the most prevalent AdV responsible for pediatric and crowded community outbreaks and was replaced by other hAdVs^16^. In addition, there is preclinical evidence that the annual immunization with the same hAdV vector may be effective due to a significant decline in vector immunity^17^. Real world evidence also shows that hAdV-specific T cell response declines with age^18^.

Moreover, the clinical data obtained with a hAdV5-based COVID-19 vaccine showed that despite the pre-existence of hAdV5-nAbs, 85%-100% of volunteers administered with only one shot of a hAd5V-based vaccine showed seroconversion against SARS-CoV-2^19^. In a two dose regimen clinical trial with the ChAdOx1-based vaccine, anti-ChAdOx1 nAbs increased with the prime vaccination but not with the boost one; that was in contrast to anti-SARS-CoV-2 nAbs that continued to increase after the boost at 28 days^20^.

We designed a novel vaccine with the aim to enhance the immunodominance of the transgene. The vaccine, named CoroVaxG.3, is based on a replication incompetent hybrid hAdV5 where the expression of a prefusion stabilized full length Spike is transcriptionally regulated by a strong promoter and an mRNA stabilizer. We also engineered the hAdV5 vector-based vaccine to display the knob domain of hAdV3 in order to improve vector targeting to muscle and dendritic cells^21,22^. Here, we describe the *in vitro* and *in vivo* data that makes CoroVaxG.3 a promising candidate to enter clinical trials.

## RESULTS

### Vaccines Construction and *in vitro* studies

In order to induce Spike immunodominance, the first step in our vaccine design was to select the most appropriate promoter to transcriptionally regulate Spike expression. Other anti-COVID 19 vaccines based on replication-incompetent AdV incorporated different alternatives of the early intermediate CMV promoter to drive Spike transcription^23–26^. However, it was shown that the CMV promoter can be silenced due to methylation, not only when it is incorporated in viral vectors that integrate to the host genome^27^, but also after *in vivo* administration of a replication-deficient adenoviral vector in the rat muscle^28^. After an extensive search in the literature, we decided to synthetize different versions of a hybrid promoter that included a CMV enhancer, the chicken β-actin promoter and a chimeric intron that contains the 5’ splice donor of the chicken β-actin 5’UTR and the 3’ splice acceptor of the minute virus of mice (MVM). This hybrid promoter was shown to drive gene expression in the rat CNS following *in vivo* transduction with AAV vectors^29^. We designed different versions that differed mainly in the size of the CMV enhancer and the β-actin promoter. In addition, we incorporated stop codons in the three open reading frames after an ATG codon 3’ splice acceptor of MVM that could interfere with Spike ATG. Two initial versions of the hybrid promoters named Pr1 and Pr2 were cloned into the pShuttle(PS)-IXP-Luc vector. Luciferase expression driven by Pr2-Luc was around 24-fold (*P* < 0.0001) higher than the control SV40 promoter, compared to only 1.6-fold increase over the SV40 promoter induced by Pr1-Luc (Fig. 1a). Based on these data, we decided to move on with Pr2 for the further constructs design.

**Figure 1.**
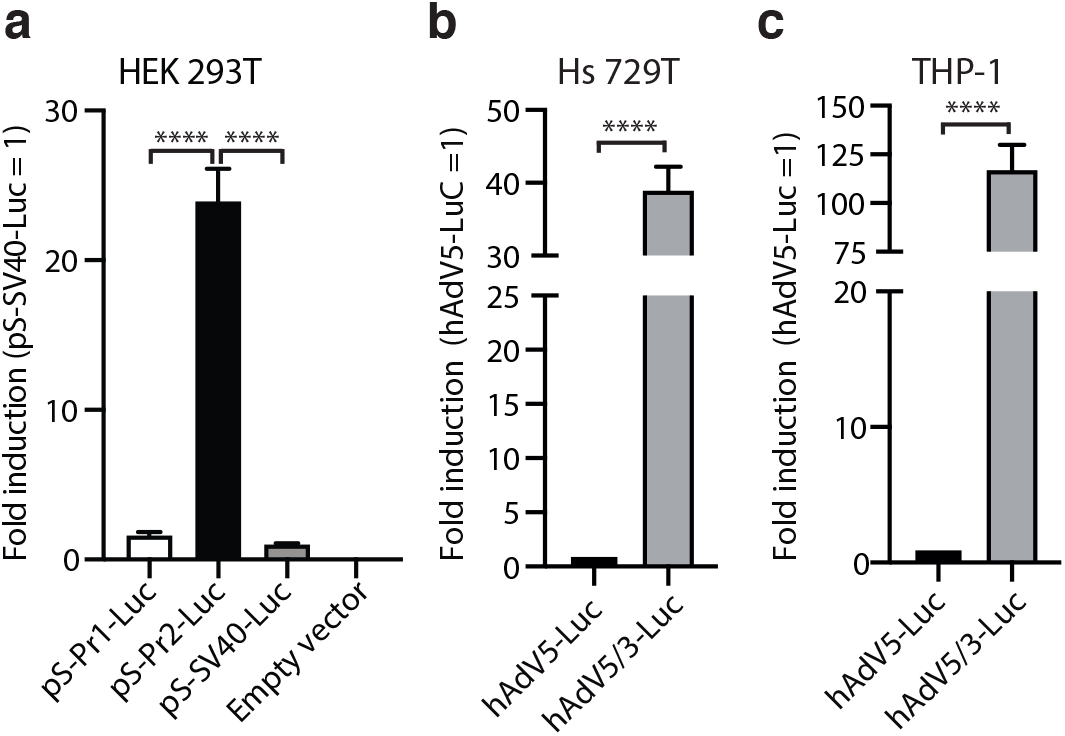
Luciferase expression following cells transfection or transduction with adenoviral vectors. **a** HEK293T cells were transiently transfected with plasmidic vectors encoding for different candidate promoters upstream of luciferase. An empty vector was included as a control. The promoter activity was evaluated by luminescence, using a luciferase gene reporter assay. The relative firefly luciferase units were nomalized by renilla and expressed as fold induction over SV40 promoter. Bars represent means ± SEM (n = 3). \*\*\*\**P* < 0.0001; one-way ANOVA. **b, c** Hs 729T and THP-1 were transduced with the replication-deficient adenoviruses, hAdV5-Luc or hAdV5/3-Luc. Luciferase activity was determined 48 hr post-transduction. Results are expressed in relative luciferase units (RLU) of Firefly Renilla normalized to hAdV5-Luc. Bars represent means ± SEM (n = 3). \*\*\*\**P* < 0.0001; Student’s t-test.

Next, we aimed to establish the adenoviral vector capacity to transduce target cells following fiber knob exchange. We initially transduced human rhabdomyosarcoma cells Hs 729T (as an example of muscle cells), and human monocytes THP-1, with replication-deficient adenoviruses hAdV5-Luc and the hybrid hAdV5.3-Luc expressing the fiber knob domain of hAdV3. Using luciferase as a surrogate marker, we observed almost 40-fold induction and more than 100-fold induction in Hs 729T and THP-1 cells, respectively, with hAdV5.3-Luc compared to hAdV5-Luc (Fig. 1b and c).

In addition to Pr2, we included the WPRE mRNA stabilizer^30^ to post-transcriptionally regulate Spike expression in our vaccines’ constructs. Among different engineered vaccine candidates expressing various Spike sequences, we selected CoroVaxG.3 for further *in vitro* and *in vivo* studies. This replication-deficient hAdV-based vaccine candidate encodes the full-length, codon-optimized sequence of SARS-CoV2 Spike protein (B.1 lineage; D614G) stabilized in its prefusion state (with the exchange of 2 prolines). CoroVaxG.3 has been additionally modified by engineering the fiber knob domain of hAdV3 instead of the knob domain of hAdV5. As a comparator we used CoroVaxG.5 that retained the native hAdV5 fiber knob domain. A replication deficient hAdV5 vector that carried no transgene was used as control (named Ad.C).

In order to establish if exchanging the fiber knob domain in CoroVaxG.3 might indeed improve muscle and dendritic cells transduction, we transduced Hs 729T cells and THP-1 monocytes with CoroVaxG.5 and CoroVaxG.3. We observed a dramatic increase in Spike expression in Hs 729T cells transduced with CoroVaxG.3 compared to CoroVaxG.5 (Fig. 2a and d). Moreover, CoroVaxG.3 was able to induce at least 6 times higher Spike expression in THP-1 monocytes compared to CoroVaxG.5 (Fig. 2b and e). Interestingly, CoroVaxG.3 induced a higher increase in Spike expression in THP-1 cells induced to differentiate to immature dendritic cells compared to undifferentiated THP-1 monocytes (Fig. 2c and f). Of note, we were unable to detect Spike expression in iDCs after cells transduction with CoroVaxG.5 even after membrane overexposure (Fig. 2e and f).

**Figure 2:**
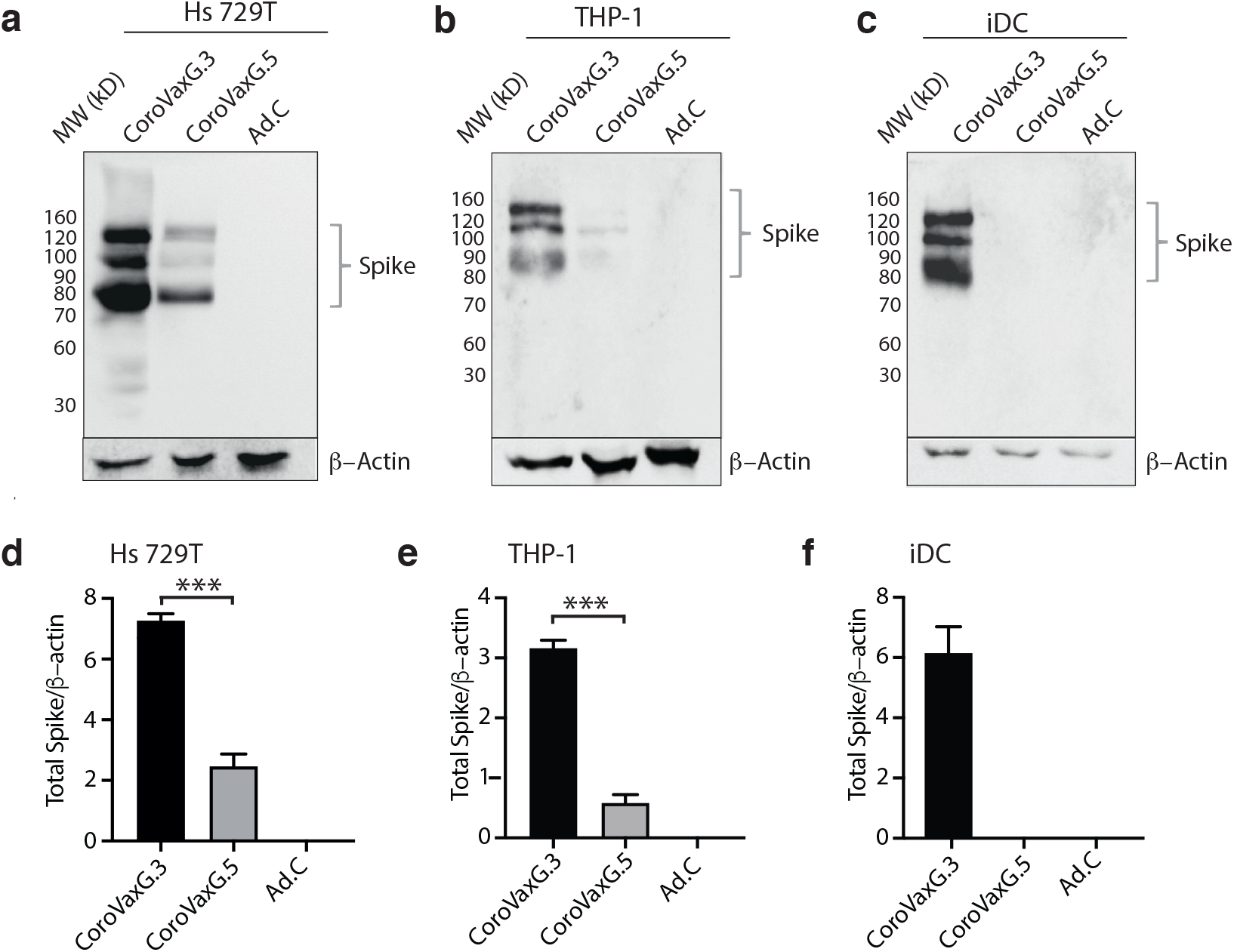
Spike expression following in vitro transduction with AdV-vectored vaccines. **a** to **c**. WB assay performed on Hs 729T, THP-1 and immature dendritic cells (iDC) cells after 48 hrs of transduction with CoroVaxG.5, CoroVaxG.3 and control Ad.C. Spike expression was detected using an anti-Spike antibody. β-actin was used as a loading control. **d** to **f** Bar graphs represent a semi-quantification of the WB assay normalized by β actin. \*\*\**P* < 0.001; one-way ANOVA.

### CoroVaxG.3 Induces Robust and Balanced Antibody Responses against SARS-CoV-2

Next, we aimed to establish whether fiber exchange might have affected the *in vivo* immune response induced by CoroVaxG.3. Thus, experimental groups of 6 to 8-week-old BALB/c mice were immunized by intramuscular injection with 10^9^ or 10^10^ viral particles (vp) of CoroVaxG.3, using as a comparator mice vaccinated with CoroVaxG.5 and with the control Ad.C at the same vp dose. Serum samples were collected at different time points after immunization (Fig. 3a), and IgG responses against Spike were evaluated by ELISA. CoroVaxG.3 induced high levels of Spike-specific IgG as early as 2 weeks after a single inoculation, very similar to the levels induced by CoroVaxG.5, while the levels induced by Ad.C were negligible (Fig. 3b, *P*< 0.05 of any vaccine vs. same dose of Ad.C at all-time points). No significant differences were observed in antibody levels between the two CoroVaxG vaccines or between doses, starting from 28 days post-vaccination. Moreover, antibody titers remained stable up to 140 days post-vaccination (maximum time point assayed) without evidence of waning. Thus, a single i.m. immunization with CoroVaxG.3 was capable of generating a robust and long-lived S-specific antibody response, with no significant differences in antibodies levels between doses. These levels were similar to those induced by CoroVaxG.5 showing that fiber exchange did not hamper the immune response induced by CoroVaxG.3.

**Figure 3.**
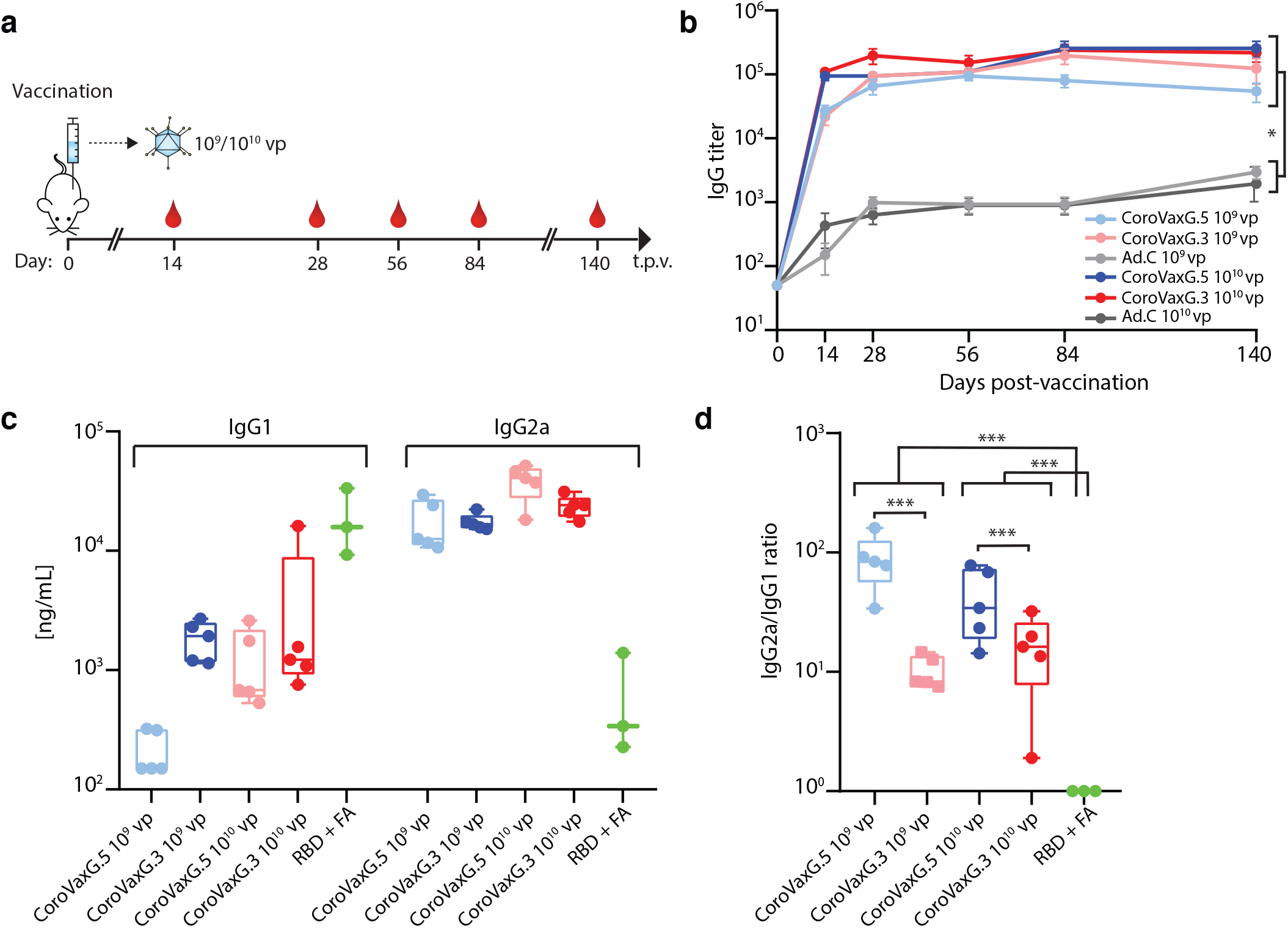
Immunogenicity and long-term humoral response induced by Ad-vectored vaccines. **a** Schedule of immunization and sample collection. Six-week-old Balb/c mice (n =5/group) received immunizations with 10^9^ or 10^10^ vp of an Ad-vectored vaccine. Sera were collected and assayed at the indicated time points (t.p.v.= time post vaccination). Animals were euthanized at 20 weeks post vaccination (long term). Serum samples were used to **b**, titer specific anti-S IgG,and **c**, **d**, to assess the IgG1 and IgG2a anti-S concentration 28 days post-vaccination. Blue symbols correspond to CoroVaxG.5; red symbols to CoroVaxG.3 and grey symbols to Ad.C; light symbols correspond to 10^9^ vp and dark symbols to 10^10^ vp. **b** Data are means ± SEM (standard error of the mean). * *P* < 0.05; two-way ANOVA with Bonferroni correction **c, d** The box plots show median, 25th and 75th percentiles and the whiskers show the range. \*\*\**P* < 0.001; one-way ANOVA with Brown-Forsythe test.

To reduce the theoretical risk of vaccine-associated enhanced respiratory disease, associated with a T_H_2-skewed response^31^ we further assessed the concentration of S-specific IgG1 and IgG2a subclasses induced by CoroVaxG.3 as a readout of T_H_1 and T_H_2 responses. As a comparator we used sera from mice vaccinated either with CoroVaxG.5 or with the SARS-CoV2-S RBD domain in Freund’s adjuvant (FA). RBD + FA vaccinated mice showed a low IgG2a/IgG1 ratio consistent with the production of high levels of IgG1 and a clear skew towards a T_H_2 phenotype (Fig. 3c and d). Interestingly, using sera samples obtained at day 28, we observed that CoroVaxG.3 and CoroVaxG.5 induced a high IgG2a/IgG1 ratio indicative of a T_H_1-biased response, although the IgG2a/IgG1 ratio was slightly higher for CoroVaxG.5 than for CoroVaxG.3 (Fig. 3d, *P* < 0.001). This difference was mainly due to a higher production of IgG1 by CoroVaxG.3, with similar levels of IgG2a induced by both vaccine candidates and doses (Fig. 3c). Similar data was obtained with sera collected 14 days after vaccination (not shown).

### CoroVaxG.3 induces a potent and durable T cell response

The generation and persistence of memory T cells provides life-long protection against pathogens and, in particular, the induction of virus-specific CD8+ T cell responses has the potential to improve the efficacy of vaccination strategies^32,33^. In order to characterize the cellular immune response induced by a single immunization with CoroVaxG.3, we assessed IFN-γ production by isolated splenocytes after specific *ex vivo* re-stimulation with Spike peptide pools, using again CoroVaxG.5 as a comparator. We observed an early strong primary immune response in vaccinated mice at 14 days post immunization, showing a similar range of IFN-γ secreting cells in both vaccinated groups (Fig. 4a). Remarkably, IFN-γ secretion was induced up to 140 days following vaccination, with no significant differences observed between the two CoroVaxG vaccines (Fig. 4b). Administration of Ad.C vector did not induce IFN-γ production at any of the assessed time points (Fig. 4a and b).

**Figure 4.**
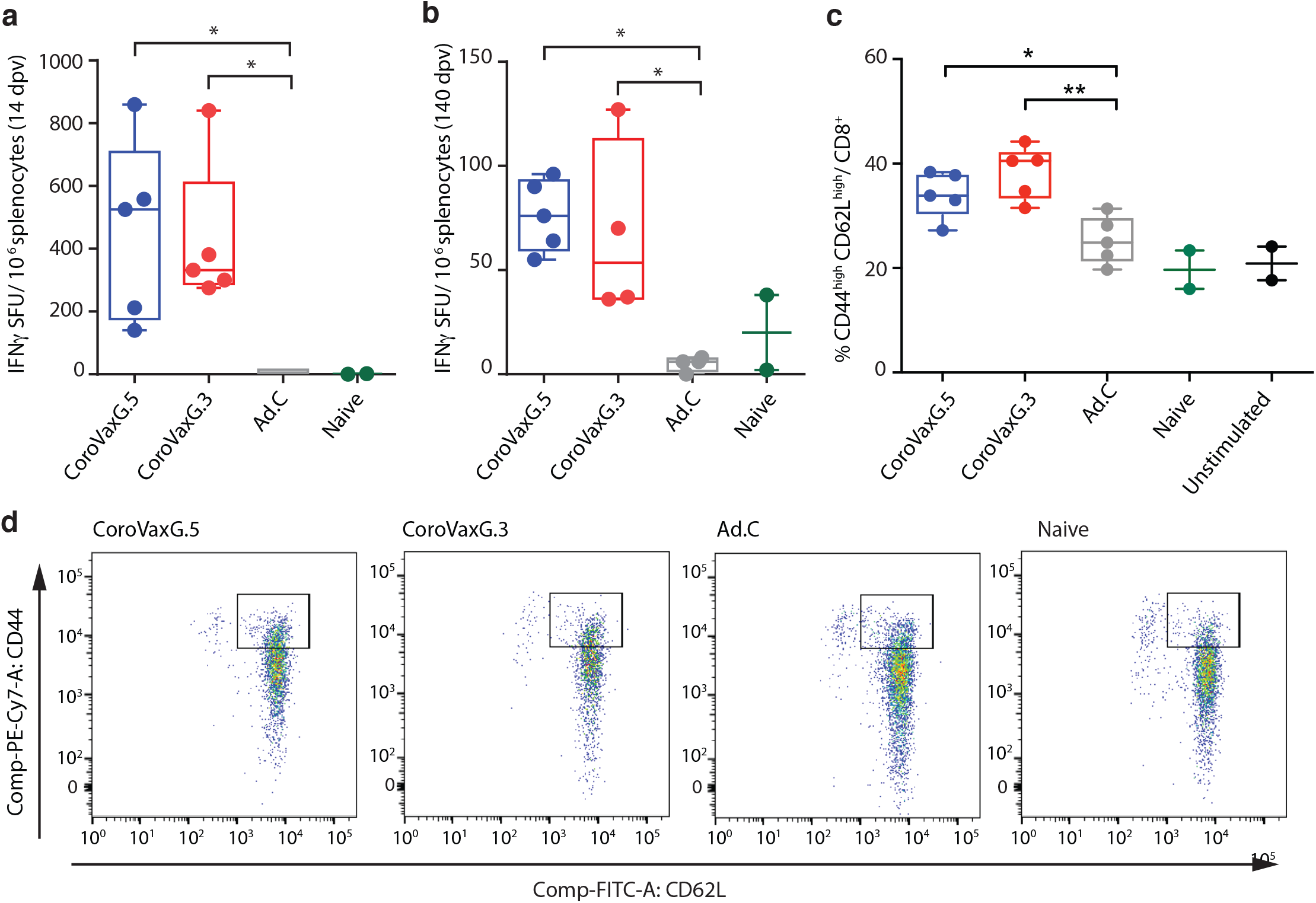
Cellular immune response induced by Ad-vectored vaccines. BALB/c mice received 10^9^ or 10^10^ vp of an Ad-vectored vaccine (blue: CoroVaxG.5; red: CoroVaxG.3; grey: Ad.C; green: Naïve) and sacrificed after 14 days (10^10^ vp) or 20 weeks (10^9^ vp). Cells secreting IFN-γ per million of splenocytes were determined by ELISPOT at **a**, 14 days and **b**, 20 weeks post immunization. Samples were analyzed in duplicates. Results of each group are expressed as mean of spot-forming units (SFU). For FACS analysis, splenocytes were stained with anti-CD8α, anti-CD62L and anti-CD44 fluorochrome-conjugated antibodies. Stained splenocytes were subjected to flow cytometry analysis to quantify memory T cells (T_CM_: CD44^high^ CD62L^high^ and T_EM_: CD44^high^ CD62L^Low^) **c** and **d**. **c** Data are expressed as percentage of total CD8^+^ cells; unstimulated controls (black dots) were included. **d** Representative dot plots of each group. The gate shows T_CM_ subpopulation. The box and whisker plots represent the median (mid-line), max and min (boxes) and range (whiskers). Vaccinated groups were always significantly different to the Naive and Unstimulated group; \**P* <0.05, \*\**P* <0.01; Kruskal-Wallis test with Dunn’s multiple comparisons *a posteriori*.

The identification of distinct memory T cells is usually based on the differential cell surface expression levels of CD44 and CD62L. Effector-memory T cells (T_EM_) located in secondary lymphoid organs can be identified as CD44^high^ CD62L^low^ while spleen located central-memory T cells (T_CM_) are identified as CD44^high^ CD62L^high 34–37^. We observed a remarkable induction of CD44^high^ CD62L^high^ CD8^+^ cells in splenocytes of vaccinated groups compared to control groups at day 140 after a single shot vaccination (Fig. 4c and d). The proportion of CD44^high^ CD62L^low^ CD8^+^ did not change in vaccinated mice at this time point (not shown). *Ex vivo* stimulated splenocytes in the Ad.C group showed levels of CD8^+^ cells expressing CD44^high^ CD62L^high^ similar to those observed in naïve mice or unstimulated splenocytes. Collectively, the data show that mice vaccinated with both vaccines developed an effective long lasting memory T-cell response against SARS-CoV-2.

### CoroVaxG.3 Induces Neutralizing Antibody Responses against VOC

To evaluate the functional quality of vaccine-generated Spike-specific antibodies, we used a pseudovirion-based neutralization assay (PBNA) to test the ability of sera from immunized mice to neutralize the entry of pseudovirus bearing Spike on their surface. Sera collected from mice at days 14, 28 and 140 after vaccination, were tested for the presence of SARS-CoV-2-specific neutralizing antibodies (nAbs) (Fig. 5a). NAbs against SARS-CoV-2 were detected in mice immunized by both CoroVaxG.3 and CoroVaxG.5. The resulting SARS-CoV-2-neutralizing activity at all-time points assayed was statistically significant (*P* < 0.0001). compared to the undetectable nAbs in the Ad.C control group, with no significant differences between the vaccines. Remarkably, this neutralizing effect was observed even after 140 days of the single dose vaccination (Fig. 5a).

**Figure 5.**
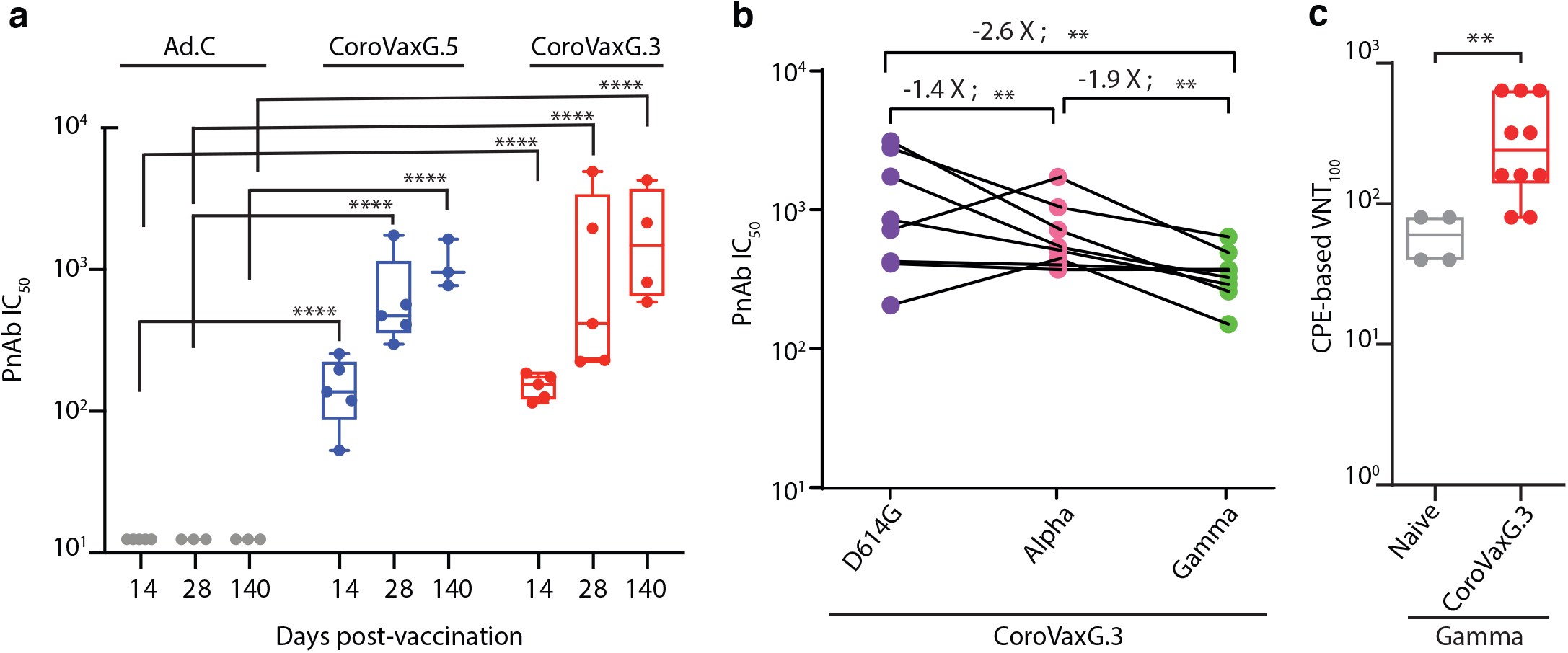
Elicitation of neutralizing antibodies by Ad-vectored vaccines. **a** Sera from animals vaccinated with the higher dose were used to measure SARS-CoV2 neutralizing antibodies by a Pseudovirus-Based Neutralization Assay. The box plots show median, 25th and 75th percentiles and the whiskers show the range. Comparisons were performed by a Kruskal-Wallis test, followed by Dunn’s multiple comparisons. **b** Changes in reciprocal serum neutralization IC_50_ values of CoroVaxG.3 vaccinated mouse sera (28 days post vaccination) against SARS-CoV-2 variants of concern. Fold change in Geometric Mean Titers relative to the WT is written above the p values. Statistical analysis was performed using a Wilcoxon matched-pairs signed rank test. Two-tailed p values are reported. **c** Neutralization against the authentic P.1/gamma VOC of CoroVaxG.3 vaccinated mouse sera (28 days post vaccination). The box plots show median, 25th and 75th percentiles and the whiskers show the range. \*\**P* < 0.01, \*\*\**P* < 0.01; two-tailed Mann-Whitney U test.

To evaluate the potential resistance of new variants to neutralization elicited by CorovaxG.3, we assessed neutralization activity against SARS-CoV-2 pseudotyped viruses containing the Spike protein of the reference B.1 strain (wild type; D614G), as well as two prevalent circulating variants in our region, B.1.1.7 (alpha, first identified in the UK) and P.1 (gamma, first identified in Manaos, Brazil). Sera from 8 mice were obtained at day 28 after vaccination with 10^10^ vp of CoroVaxG.3 and sera from 4 naïve mice were used as a control. All naïve sera showed undetectable levels of neutralization against the tested SARS-CoV-2 variants (IC_50_<25). The neutralizing antibody titers elicited against the wild-type strain (GMT = 870.7, CI 95% 384.8–1970) showed a slight decrease (1.4 fold, *P*< 0.01) versus B.1.117 lineage (GMT = 627.1, CI 95% 402.9–976.2) and a larger but still moderate decrease (2.6 fold, *P*< 0.01) versus P.1 lineage (GMT = 334, CI 95% 232.5–480) (Fig. 5b). Interestingly, all sera tested neutralized the pseudovirus variants with IC_50_s of at least 150. Assessment of the same sera samples from day 28 used for the PBNA studies, confirmed that CoroVaxG.3 was able to elicit neutralizing antibodies against the authentic P.1/gamma VOC (Fig. 5c).

## DISCUSSION

In the present study we provide evidence on a novel, replication-deficient hAdV-based vaccine, aimed at achieving Spike immunodominance to skew the immune response towards the transgene, and reducing the vector-targeted immune response that will occur despite the vector used. Immunodominance of Spike was tackled from different angles: (i) optimizing Spike expression using a potent promoter and an mRNA stabilizer; (ii) stabilizing Spike in a prefusion conformation to enhance Spike immunogenicity^38,39^; (iii) engineering the hAdV5 fiber to express the fiber knob domain of hAdV3 to enhance transduction of muscle and dendritic cells.

We observed a dramatic enhancement of Spike expression in human muscle cells as well as in human monocytes and dendritic cells with CoroVaxG.3 compared to CoroVaxG.5. Since in both vaccine candidates Spike expression was under the regulation of Pr2 and the WPRE mRNA stabilizer, we concluded that the enhanced Spike expression in human muscle derived cells, monocytes and dendritic cells was due to the fiber knob exchange as demonstrated also through the studies using replication-deficient vectors expressing luciferase.

CoroVaxG.3 was able to induce strong humoral and cellular immune responses similar to the levels previously reported in other preclinical animal studies, including the production of anti-SARS-CoV2 neutralizing antibodies^38–40^. Despite the significant difference in Spike expression in the human targeted cells *in vitro*, the *in vivo* potency of both vaccines in mice was quite similar, although CoroVaxG.3 induced higher levels of IgG1 (see below). A likely explanation is that CoroVaxG.3 uses alternative receptors in mice to those used to transduce human cells. Indeed, murine DSG-2 is not recognized by hAdV3 or vectors derived from this hAdV serotype^41^; thus, murine cells transduction *in vivo* likely occurred through alternative receptors. Moreover, although hAdV expressing the serotype 3 fiber were shown to efficiently transduce human monocytes and skin resident DC (Langerhans cells and skin DC), transduction occurred through binding to the B7-family members CD80/CD86 without affecting subsequent T cell stimulation^42^, suggesting that CoroVaxG.3 can transduce target cells through alternative receptors. In this regard, it has been shown that the development of protective cellular immunity induced by adenoviral-based vaccines correlated with the magnitude and persistence of the transgene expression^43^. Thus, even if we were unable to see remarkable differences with CoroVaxG.3 and G.5 vaccines in the immune response in the mice model, it is likely that a persistently high Spike expression in vaccinated individuals following immunization with CoroVaxG.3, would enable the induction of a sustained and enriched immune response.

One of the main initial concerns regarding vaccine development was the possibility of vaccine-related diseases, either induced by antibodies (antibody-dependent enhanced disease, ADE), which is associated with the presence of non-neutralizing antibodies, or vaccine-associated enhanced respiratory disease (VAERD), linked to inflammation induced by Th2-skewed immune responses^44,45^. On the other hand, the induction of an unbalanced inflammatory Th1-biased response by some Ad vectors, due to an exacerbated production of type I interferons, has been suggested to impair transgene expression levels, dampening subsequent humoral immune responses^46^. Therefore, the induction of a balanced immune response is desirable. Interestingly and despite the fact that both vaccines candidates induced a robust and long lasting humoral and cellular immune response, CoroVaxG.3 elicited higher levels of IgG1 and hence a more balanced Th1/Th2 ratio, evidencing an optimal immune profile in mice. The difference in the subclass profile between the two vaccines might be related to the differential tropism observed in the *in vitro* studies. It was already shown that mice injected with DCs transduced *ex vivo* with hAdV5 expressing βgal elicited mainly IgG2a antibodies, while vaccination with hAdV5-βgal transduced myoblasts elicited a more balanced Ab response with an IgG1/IgG2a ratio similar to that induced by direct vaccination with hAdV5-βgal ^47^; thus, it is likely that CoroVaxG.3 is inducing a more balanced immune response due to its enhanced tropism for muscle cells.

In acute and convalescent COVID-19 patients it has been observed that the presence of T cell responses is associated with reduced disease^48–50^, suggesting that SARS-CoV-2-specific T cell responses may be important for control and resolution of primary SARS-CoV-2 infection^51^. Preclinical studies in macaques have shown that both neutralizing antibody titers and Fc functional antibody responses correlated with protection^52,53^ and that purified IgG from convalescent animals, in the absence of cellular and innate immunity, effectively protected naïve recipients against a challenge with SARS-CoV-2^54^. However, in the setting of waning and subprotective antibody titers, cellular immune responses were critical for rapid virological control. Through CD8 depletion studies, it was shown that cellular immunity, especially CD8+ T cells, contributed to protection against rechallenge with SARS-CoV-2^51^. The present data show that CoroVaxG vaccines induced a rapid and long lasting cellular immune response, as characterized by IFN-γ production and proliferation of memory CD8+ T cells, upon re-stimulation with SARS-CoV-2 peptides. *Ex vivo* stimulated splenocytes obtained from naïve mice, or from mice injected with the empty vector, and non-stimulated splenocytes obtained from vaccinated mice exhibited basal levels of non-specific CD8+ memory cells consistent with aged mice^55,56^. Roberts *et al* ^57^ showed that both CD62L^low^ and CD62L^high^ memory cell subpopulations contributed to recall responses to Sendai virus infection, however, the relative contributions of these subpopulations changed over time: CD62L^low^ cells dominated at early time points, whereas CD62L^high^ cells dominated at later time points. Consistent with this, we observed that *ex vivo* stimulation of vaccinated mice splenocytes with Spike peptides induced a long term increase in the T_CM_ subpopulation.

A correlate of protection that allows evaluation of vaccine efficacy based on immune readouts has been sought since the beginning of COVID-19 vaccine development. Recent studies suggest that neutralizing antibodies could serve this purpose and are, therefore, the main parameter that might help to predict the success of a vaccine candidate^58^. CoroVaxG.3 was able to elicit neutralizing antibodies to a similar extent to the levels previously reported in preclinical animal models, and those nAbs were stable in the long term. Moreover, studies with pseudoviruses and authentic SARS-CoV-2 containing variant substitutions suggested that neutralizing ability of the antibodies in sera raised after CoroVaxG.3 vaccination is only slightly reduced but overall largely preserved against the B.1.1.7 and P.1 lineages that are VOC prevalent in Latin America^59^. This mild reduction in neutralization by vaccine-raised sera against these VOC is in the same range than that observed for sera of patients who received two doses of either the BNT162b2 Pfizer-BioNTech or ChAdOx1 nCoV-19 Oxford-AstraZeneca vaccine^60^.

Each one of the AdV-based vaccines already approved by different regulatory bodies and the CoroVaxG.3 platform described in this study are based on different AdV species or human serotypes, are designed in a different way and most importantly, bind to different cell surface receptors^10^. Although some adenoviral vectors from human and non-human primate origin display robust immunogenicity *in vivo* comparable to that of AdV5, they are less immunogenic^61,62^, and there are considerable differences in the phenotype and functionality of the immune response they can elicit^15,43,61^. These differences could clearly impact on the durability, potency and quality of the immune response induced by the different adenoviral-based vaccines. Thus, it is likely to assume that they might trigger a differential immune response that will not necessarily be visible during the short time that evolved since immunization with the different vaccines started, and might require an extended follow-up period to emerge. How the different immune profiles will influence vaccine efficacy in patients remains to be seen. We developed a novel anti-COVID-19 vaccine based on a hybrid hAdV5 vector that expresses a chimeric fiber and where the high expression of Spike stabilized in its prefusion state is supportive to achieve its *in vivo* immunodominance. CoroVaxG.3 showed differential characteristics compared to CoroVaxG.5, a vaccine comparator with features similar to many vaccines that gained regulatory approval. Our study demonstrated that the high levels of neutralizing antibodies elicited by CoroVaxG.3 were maintained for at least 5 months, and probably much longer, since they were stable over the analyzed period. Moreover, sera obtained from CoroVaxG.3 vaccinated mice, were able to neutralize VOC of regional importance. An anti-COVID 19 vaccine based on the molecular design of CoroVaxG.3 is ready to enter clinical trials in the next coming months based on a one shot administration.

## METHODS

### Reagents and Cells

HEK293T (CRL-3216), Vero cells (CCL-81), Hs 729T (HTB-153) and THP-1 (TIB-202) cells were obtained from the ATCC (Manassas, VA, USA). HEK293 cells were purchased from Microbix Biosystems Inc (Mississauga, Canada) and 911 cells were already described^63^. HEK293T-hACE2 cells were already described^64^. All the cell lines were grown in the recommended medium supplemented with 15% of fetal bovine serum (Natocor, Cordoba, Argentina), 2 mM glutamine, 100 U/ml penicillin and 100 μg/ml streptomycin and maintained in a 37 °C atmosphere containing 5% CO_2_. For HEK293T and HEK293T-hACE2 cell cultures non-essential amino acids (1X final concentration) were added. Immature dendritic cells (iDC) were generated from THP-1 monocytes as previously described^65^. To induce differentiation, THP-1 monocytes were cultured during 5 days in RPMI-1640 Medium (Gibco, MD, USA); 2-mercaptoethanol (0.05 mM final concentration; Gibco, MD, USA) and fetal bovine serum (10%); adding rhIL-4 (100 ng = 1500 IU/ml; Peprotech, NJ, USA) and rhGM-CSF (100 ng = 1500 IU/ml; Peprotech, NJ, USA). The acquired properties of iDCs were analyzed under microscope. Medium exchange was performed every 2 days with fresh cytokine-supplemented medium.

### Promoter and fiber selection

Promoters were synthesized by Genscript (NJ, USA) with *Not*I / (*Xho*I-*Stu*I) flanking restriction sites and cloned in the *Not*I / *Stu*I sites of the vector pShuttle-I-XP-Luc^63^ to obtain pShuttle-Pr1-Luc and pShuttle-Pr2-Luc. The vectors pAd-SV40-Luc, and pS-CMV-Renilla were previously described^63^. For the selection of the most appropriate promoter, HEK293T cells grown in 24-well plates were co-transfected with 1 μg of the different plasmids and 100 ng of pS-CMV-Renilla, using Lipofectamine 2000 (Thermo Fisher Scientific, CA, USA). Twenty-four hours later, the cells were collected and assayed for Firefly and Renilla Luciferase activities using the Dual-Luciferase Reporter Assay System (Promega, WI, USA) and measured in a Genius luminometer (TECAN, Maennedorf, Switzerland). Each experiment was performed at least three times. hAdV5/3-Luc and hAdV5-Luc replication-deficient adenoviral vectors were already described^63^. Hs 729T and THP-1 cells were transduced with hAdV5-Luc and hAdV5/3-Luc viruses at MOI 500. Forty-eight hours later, the cells were collected and assayed for Renilla Luciferase activity as described.

### Vaccine design and production

The sequence of the Spike protein gene was extracted from the official GISAID reference sequence WIV04 (https://www.gisaid.org/) and modified to obtain the D614G, K986P and V987P variant named D614G-PP. A cloning cassette flanked by *Stu*I / *Sal*I restriction sites was synthesized by Genscript (NJ, USA) including a Kozak consensus sequence (GCCACCATG), the codon optimized Spike (D614G–PP), 589 bp of the woodchuck hepatitis virus posttranscriptional regulatory element (WPRE)^30^ and 222 bp of the SV40 virus late polyadenylation signal. Codon optimization was performed with the Vector Builder software (https://en.vectorbuilder.com/tool/codon-optimization.html). The synthesized 4,650 bp fragment was cloned into the pShuttle-Pr2-Luc vector digested with *Stu*I / *Sal*I to exchange the luciferase ORF by the designed Spike cassette, downstream of Pr2. The sequence of the resulting plasmid pS-Spike(D614G)-PP was confirmed by sequencing (Macrogen, Seul, Korea). To construct the non-replicating adenoviruses, the plasmid pS-Spike(D614G)-PP was linearized with *Pme*I and co-transformed with E1/E3 (pCoroVaxG.5) or E1 (pCoroVaxG.3) deleted adenoviral backbone vectors in electrocompetent BJ5183 bacteria. The identity of the plasmids was confirmed by sequencing. The recombinant DNAs were linearized with *Pac*I and transfected into 911 cells. The viruses were propagated in HEK-293 cells in CellSTACK^®^ cell culture chambers (Corning, Arizona, USA), purified by double CsCl density gradient centrifugation and stored in 10% glycerol in single-use aliquots at –80°C.

### Western blots

To assess Spike expression by western blot, 1 × 10^6^ cells were seeded and cultured in 6-well plates. THP-1 (human monocytes), iDC and Hs 729T (rhabdomyosarcoma) cells were transduced with CoroVaxG.5, CoroVaxG.3 or Ad.C (MOI of 1000 for THP-1 and 500 for Hs 729T). Cells were washed twice with ice-cold PBS and lysed in Laemmli sample buffer 2X. Protein extracts were separated by SDS-PAGE with a 10% gel and transferred to nitrocellulose membranes (Bio-Rad Laboratories). The membranes were probed with anti-spike Ab (40150-T62, Sino Biological) and anti-beta-actin Ab (A4700; Sigma). After incubation with HRP-AffiniPure Goat Anti-Rabbit IgG (Jackson ImmunoResearch), chemiluminiscence was detected with ECL following the manufacturer’s instructions (Amersham)and digitized by Image Quant LAS 4000 (GE-Cytiva MA, USA). Semi-quantifications of WB assays were performed by densitometry using the ImageJ software 1.53 (Wayne Rasband, NIH, USA), normalizing by β-actin expression.

### Mice immunization

Six- to eight-week-old male BALB/c mice (obtained from the animal facility of the Veterinary School, University of La Plata, Argentina) were immunized with 10^9^ or 10^10^ viral particles (vp) of Ad.C (empty vector), CoroVaxG.5 or CoroVaxG.3 in 30 μl PBS via intramuscular injection in the hind leg. Serum samples for intermediate time points were obtained by submandibular bleeds for humoral immune response analyses. Final serum samples were obtained via cardiac puncture of anesthetized mice. The collected whole blood was allowed to clot at 37 °C for 1 h before spinning down at 500 × g for 10 min. The clarified sera were stored at −20 °C. For RBD inoculations eight-week-old male BALB/c mice were immunized with 7.5 μg of the receptor binding domain of Spike protein (kindly gifted by Andrea Gamarnik, Argentina) in 75 μL Complete Freund’s Adjuvant (CFA, Sigma, St. Louis, MO) via subcutaneous injection and boosted 2 weeks later with 5 μg of RBD in 100 μL Incomplete Freund’s Adjuvant (IFA, Sigma). Mice were bled 14 days after the boost. Mice were maintained under specific pathogen-free conditions at the Institute Leloir animal facility and all experiments were conducted in accordance with animal use guidelines and protocols approved by the Institutional Animal Care and Use Committee (IACUC protocol 69).

### ELISA

Sera from all mice were collected at different time points after immunization and evaluated for SARS-CoV-2-S-specific IgG antibodies using ELISA. Sera collected at week 4 after vaccination were also tested for SARS-CoV-2-S-specific IgG1 and IgG2a antibodies using ELISA. Briefly, ELISA plates (BRANDplates^®^, immunoGrade, BRAND GMBH + CO KG) were coated with 100 ng of recombinant SARS-CoV-2 Spike protein (S1+S2 ECD, His-tag, Sino Biological) per well overnight at 4 °C in 50 μL PBS and then blocked with PBS-T / 3% BSA (blocking buffer) for one hour. Plates were subsequently incubated 1 h at room temperature with three-fold dilutions of the mouse sera in blocking buffer. Plates were washed and bound specific IgG was detected with a HRP-conjugated goat anti-mouse IgG H&L antibody (ab6789, Abcam) diluted 1: 10,000 in blocking buffer. Color development was performed by addition of 50 μL TMB Single Solution (Thermo Fisher Scientific). After 8 min, the enzyme reaction was stopped with 50 μL of 1 M sulfuric acid per well and the absorbance was measured in Bio-Rad Model 550 microplate reader (Bio-Rad Laboratories). Sera were assayed in duplicates and antibody titer represents the last reciprocal serum dilution above blank.

### ELISA for quantification of IgG subclasses

For IgG1 and IgG2a ELISAs, plates were coated with SARS-CoV-2 Spike protein as described in the previous section. The S-specific IgG1e3 and IgG2a mAbs (Invivogen) were serially diluted from 200 ng/mL to 3.125 ng/mL in blocking buffer and incubated 1 h at room temperature. Mouse sera were diluted 1:150 or 1:1500 in blocking buffer in order to fit the linear range of the standard curve. After the plates were washed, HRP-conjugated goat anti-mouse IgG1 and IgG2a (1:20000, ab97240 and ab97245, Abcam) were added to each well and the ELISA was performed as described before. For each IgG subclass reference, a standard curve was plotted using GraphPad Prism 8.0 generating a four-parameter logistical (4PL) fit of the OD 450 nm at each serial antibody dilution. In this way, the relative levels were comparable between IgG subclasses measured on the same antigen. As a comparator, the sera from 3 mice inoculated with RBD + Freund’s adjuvant was included.

### Pseudovirus construction for *in vitro* neutralization assays

The pseudoviral particles (PVs) containing SARS-CoV2 Spike-D614G protein were generated according to the methodology described by Nie et al ^66^, with modifications. Basically, we generated a replication defective Vesicular Stomatitis Virus (VSV) PV in which the backbone was provided by a pseudotyped ΔG-luciferase (G*ΔG-luciferase) rVSV (Kerafast, Boston, MA, USA), that packages the expression cassette for firefly luciferase instead of VSV-G in the VSV genome. Briefly, the full length cDNA of Spike-D614G, and the Spike variants B.1.1.7 (alpha, first identified in the UK) and P.1 (gamma, first identified in Manaos, Brazil), were cloned into the eukaryotic expression vector pcDNA3.1, using the EcoRV restriction site and blunt-end ligation strategy, to generate the recombinant plasmids pcDNA-3.1-Spike-D614G, pcDNA-3.1-Spike-B,1,1,7 and pcDNA-3.1-Spike-P.1.. HEK-293T cells growing in Optimem media (Gibco, MD, USA) with 2% of FBS were transduced with G*ΔG-VSV at a multiplicity of infection of four. Twenty-minutes later, the cells were transfected with 30 μg of pcDNA-3.1-Spike-D614G, using Lipofectamine 3000 (Thermo Fisher Scientific, CA, USA) and incubated six hours at 37°C, 5% CO_2_. Then, the cells were washed four times with PBS in order to remove all the residual G*ΔG-VSV, and cultured in complete media at 37°C, 5% CO2. After 48 hours the supernatant containing the PVs was collected, filtered (0.45-μm pore size, Millipore) and stored in single-use aliquots at −80 °C. The 50% tissue culture infectious dose (TCID_50_) of SARS-CoV-2 PV was determined in sextuplicates and calculated using the Reed–Muench method as previously described ^66^.

### Pseudovirus Based Neutralization Assay

The neutralization assays were performed as previously described^66^ with some modifications. Briefly, 50 μl of serially diluted mouse sera were combined with 65 TCID_50_ PVs in 50 μl of complete medium (DMEM supplemented with 10% FBS and non-essential aminoacids) in 96 well plates (Greiner Bio-One, Germany) and incubated at 37°C, 5% CO_2_ for 1 h. Next, 100 μl of 5×10^5^/mL HEK293T-ACE2 cells were added to the pseudovirus-serum mixture and incubated at 37 °C, 5% CO2 for 20-24 h. Conditions were tested in duplicate wells on each plate and a virus control (VC = no sera) and cell control (CC = no PV) were included on each plate in six wells each to determine the value for 0% and 100% neutralization, respectively. Media was then aspirated from cells and Firefly luciferase activity was determined with the Luciferase Assay System (Promega) as recommended by the manufacturer. The percentage of inhibition of infection for each dilution of the sample is calculated according to the RLU values as follows: % inhibition = [1 – (average RLU of sample – average RLU of CC) / (average RLU of VC – average RLU of CC)] × 100%. On the basis of these results, the IC50 of each sample was calculated by the Reed-Muench method^66^.

### Neutralization of authentic SARS-CoV-2 virus

Neutralizing antibody (nAb) titers against SARS-CoV2 were defined according to the following protocol. Briefly, 50 μl of serum serially diluted two-fold from 1:5 to 1:640 were added in duplicates to a flat bottom tissue culture microtiter plate (COSTAR^®^ 96 well plates), mixed with an equal volume of 110 PFU of a SARS-CoV-2 Gama isolate (P.1, CD1739-P4/2020, GenBank Accession number: MZ264787.1, May, 2021, kindly gifted by Dr. Esther C. Sabino from Instituto de Medicina Tropical (USP, São Paulo-SP, Brazil). All dilutions were made in MEM with the addition of 2.5% Fetal Bovine Serum (ThermoFisher Scientific) and the plates were incubated at 37 °C in 5% CO_2_. After 1 h incubation the virus-sera mixtures were added to each well of a flat bottom tissue culture microtiter plate (Greiner CELLSTAR^®^ 96 well plates) containing 5×10^4^ Vero cells per well (90% of confluency). After 72 h of incubation VNT was evaluated by optical microscopy of the cell culture. Neutralizing titer was the last dilution in which we did not observe a cytopathic effect (CPE). A positive titer was equal or greater than 1:10. Sera from naïve mice were included as a negative control.

### IFN-γ ELISPOT

Spleens were removed from vaccinated or control BALB/c mice at 14 and 140 days post immunization and splenocytes were isolated by disaggregation through a metallic mesh. After RBC lysis (Biolegend), resuspension, and counting, the cells were ready for analysis. The IFN-γ-secreting cells were assessed using the ELISPOT mouse IFN-γ kit (R&D Systems) according to the manufacturer’s protocol. Cells were cultured for 18 h at 5×10^5^ cells per well with 2 μg/ml of a peptide pool consisting mainly of 15-mers (overlapping by 11 amino acids) covering the immunodominant sequence domains of the SARS-CoV-2 Spike protein (PepTivator^®^ SARS-CoV-2 Prot_S; Miltenyi Biotec, Germany). The number of spots was determined using an automatic ELISPOT reader and image analysis software (CTL-ImmunoSpot^®^ S6 Micro Analyzer, Cellular Technology Limited (CTL), Cleveland, USA).

### Flow Cytometry

Splenocytes from vaccinated or control BALB/c mice at 20 weeks post immunization were obtained as mentioned before. The cells were incubated for 18 h at 37 °C at 1.5×10^6^ cells per well with 2 μg/ml of SARS-CoV-2 Spike protein peptide pool (PepTivator^®^ SARS-CoV-2 Prot_S, Miltenyi Biotec, Germany). Cells were stained with anti-CD8α (APC), anti-CD62L (FITC) and anti-CD44 (PE Cy7) surface markers (Biolegend). Cells were acquired on a FACSAria Fussion cytometer and analysis was performed using Flow Jo version 10.7.1.

### Statistical Analysis

All quantitative data are presented as the means ± SEM. ANOVA and two-sample t tests were used to compare continuous outcomes between groups. Differences were considered significant if P < 0.05. The statistical tests used are indicated in each figure legend. For S-specific binding antibodies as measured by ELISA (Fig. 3b) and PnAb titers as measured by PBNA (Fig. 5a), data were log-10 transformed prior to statistical analysis. Statistical differences between immunization regimens and time points after immunization were evaluated two-sided using a two-way ANOVA and a multiple comparisons Bonferroni correction was performed *a posteriori*. To compare isotype ratios between treatments data were log-10 transformed and a one-way ANOVA Brown-Forsythe test was applied. The change in neutralization for the different VOC was assessed by a Wilcoxon matched-pairs signed rank test. Comparison between the vaccinated and naïve sera in the VNT was done applying a two-tailed Mann-Whitney U test. All analyses were conducted using GraphPad Prism software (version 8.2). All statistical tests were conducted two-sided at an overall significance level of α = 0.05.

## ACKNOWLEDGMENTS

We thank Andrea Gamarnik and María Mora Gonzalez Lopez Ledesma for kindly providing samples of Spike RBD. This study has been supported in part through a Sponsored Research Agreement of Fundacion Instituto Leloir-CONICET(IIBBA) with Vaxinz

## AUTHOR CONTRIBUTIONS

Design of vaccines: M.V.L., E.G.A.C., S.E.V., F.J.N., A.S.L, O.L.P.

Designed studies and analyzed data: M.V.L., S.E.V., E.G.A.C., F.J.N., A.S., P.M.B., O.L.P.

Performed experiments: M.V.L., S.E.V., E.G.A.C., F.J.N., A.S., P.M.B., M.S-L., J.A., D.A-C., G.D.R., J.T.M., C.T.B., V.B.D., T.M.D., T.C.D.B., L.M.R.G.

Contributed resources: K.G., H.O., M.J.B.C.G

Drafted and revised the manuscript: O.L.P., S.E.V., P.M.B., M.V.L., F.J.N., E.G.A.C., J.T.M.

